# Cardiac interoception impacts behavior and brain-wide neuronal dynamics

**DOI:** 10.1101/2025.04.25.650721

**Authors:** Kristian J Herrera, Arman Zarghani-Shiraz, Misha B Ahrens, Florian Engert, Mark C Fishman

## Abstract

We sought to explore the question as to whether an animal’s behavior can be modified by internal physiological changes. We focused on the optomotor response, in which an animal moves in response to visual gratings, because it is quantitative, robust, and evolutionarily conserved. Using larval zebrafish, we demonstrate that engagement in the optomotor response is inversely related to heart rate. We modulate heart rate by external threat, activation or blockade of the sympathetic nervous system, pharmacological blockade of the cardiac pacemaker channel, and direct optogenetic pacing of the heart, and find that the correlation persists through all perturbations. We find neurons in the primary sensory ganglia and several regions of the brain whose activity reflects changes in heart rate, some more active during bradycardia and some during tachycardia. Specifically, we show that the area postrema, known to be a center of cardiovascular integration, shows particularly strong encoding of heart rate, both following threat and during optogenetic cardiac pacing. We suggest that there may be neural mechanisms to assess heart rate changes over time, and that this interoceptive measurement is used to regulate other neural circuits and behavioral output.

## INTRODUCTION

The nervous system of all vertebrates is responsible for monitoring and regulating the internal state of the body. This process, collectively known as interoception, is crucial for preserving bodily homeostasis^1^ and relies on the coordinated action of the autonomic and central nervous systems. Broadly speaking, there are two types of interoceptors, those that sense chemical changes, termed chemoreceptors, and those that sense stretch, termed mechanoreceptors^2^. Many of the primary interoceptive sensory neurons reside in two ganglia, the glossopharyngeal and nodose (vagal) ganglia. These neurons receive signals from internal organs using sensory axons and propagate this information primarily to brainstem neurons in the area postrema and the nucleus tractus solitarius. These cells, in turn, make both local connections, as well as long-range connections widely throughout the brain. Such local connections play an important role in guiding critical homeostatic reflexes, such as the baroreflex^3^, Bezold-Jarisch reflex^4^, and emesis^5^. This is largely accomplished by regulating the activity of the key efferent control systems for the viscera, the sympathetic and parasympathetic limbs of the autonomic nervous system. Less clear is what role ascending projections to the rest of the brain may play.

More than one hundred years ago, Lange and James proposed that emotions are a consequence of physiological state, such that, for example, fear reflects the increase in heart rate^6,7^. Recent evidence has supported this notion, with anxiety-like behaviors enhanced in mice with optogenetically elevated heart rates^8^. Clinical research offers evidence that maladies of the autonomic nervous system are associated with neuropsychiatric conditions, including autism spectrum^9^, schizophrenia^10–12^, and manic depressive disorders. This is mirrored by the thera- peutic promise of vagal nerve stimulation therapy, which, through still unknown mechanisms^13^, has aided patients afflicted with drug-resistant epilepsy^14–17^ and manic depressive disorder^18^. These observations suggest that interoception may modulate not only emotional state but behaviors more generally. To explore this hypothesis, we used the larval zebrafish, where, in awake and behaving animals, we can use non-invasive, entirely optical methods to quantify visceral dynamics, while monitoring and manipulating activity of individual neurons of the peripheral ganglia and central nervous system. We focused on whether and how changes in cardiovascular physiology might affect the robust optomotor response, where the fish is swimming and turning to follow whole field motion stimuli. Over the past decade, biologically plausible circuits have been proposed to explain how motion is initially detected in the retina and its midbrain arborizations fields^19–22^, and subsequently integrated over space^22^ and time to promote accurate behavioral responses^23,24^. Plasticity in the context of feedback control has been explored extensively in this system^25–27^, and specific models of neural and cellular implementations have started to emerge. Thus, by looking at how visceral information may affect this well-characterized neural transformation, we are primed for a detailed circuit dissection at the systems and cellular level, where the small size and optical transparency of the larval zebrafish can be leveraged.

Here we find that the optomotor response (OMR) varies with heart rate and that this relationship holds true regardless of whether heart rate changes are spontaneous, linked to struggle, driven pharmacologically or by neural stimulation, or changed by direct optogenetic pacing of the heart. This suggested that there might be neural systems in place to assess these longer term changes in heart rate. We indeed identified cells in the nodose ganglia and in specific regions of the central nervous system whose activity was either correlated or anti-correlated with heart rate. In one central nervous system region in particular, the area postrema, known to be a center for cardiovascular integration, we found that direct manipulation of heart rate could trigger calcium responses. Thus, in addition to input from baroreceptors and oxygen chemoreceptors, the cardiovascular interoceptive input to the brain appears to contain cells and circuits capable of integrating the heart rate over time, and utilizing such heart rate input to modulate motor behavior.

## RESULTS

### Zebrafish alter cardiodynamics following threats

To study how changes in cardiodynamics and behavioral state are coupled in larval zebrafish, we aimed first to establish a paradigm for eliciting and recording robust changes in heart rate while tracking motion of the tail^28^. With this system, we could capture heart and tail motion through a single camera placed below the animal (Figure 1A). In selecting stimuli, we reasoned that cues leading to high-energy behaviors, such as struggles, are likely to induce changes of internal state and spur changes in heart rate^29^. Thus, we tested two stimuli, a visual “dark flash” and a chemical “salt pulse” that elicit struggles in head-embedded larvae^28,30^ (Figure S1A).

**Figure 1.**
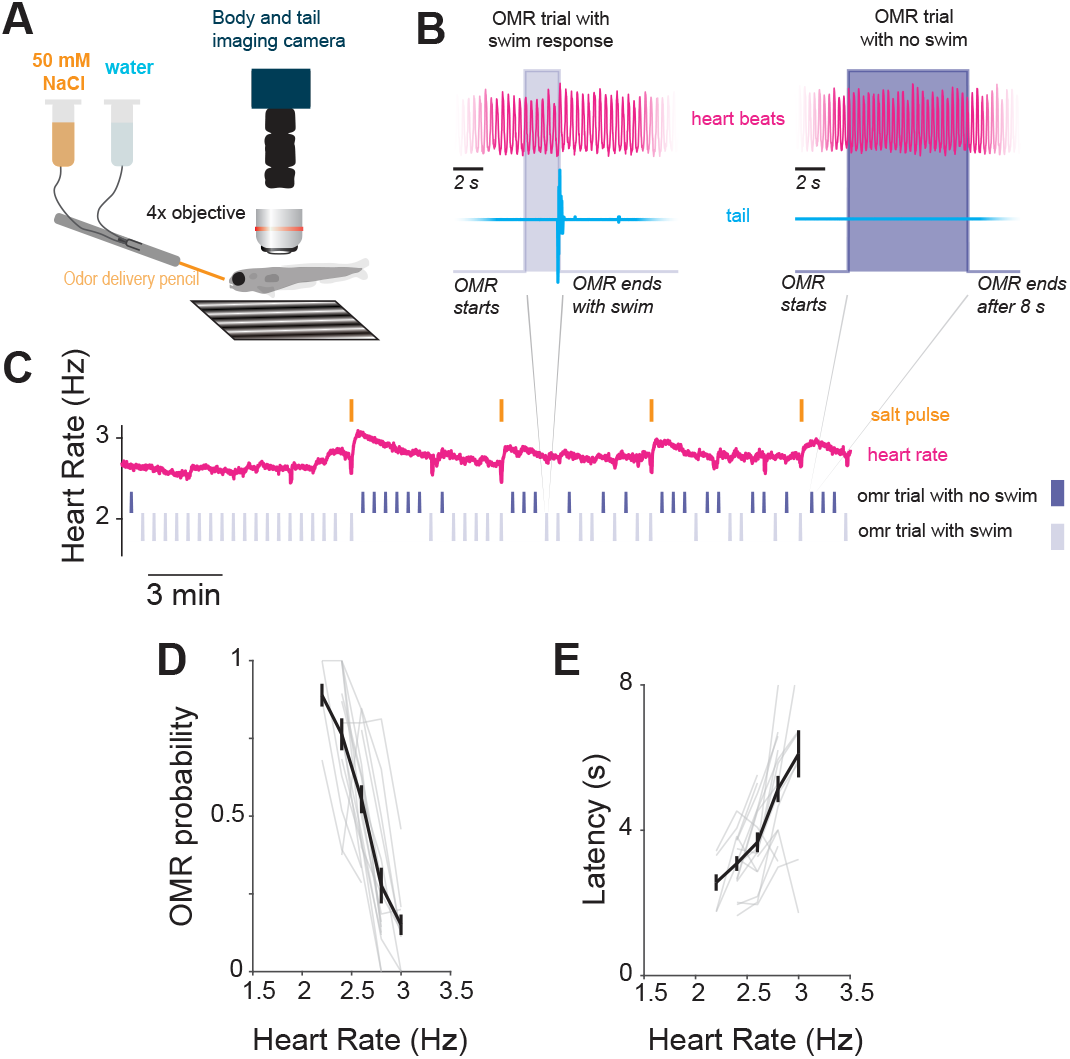
Optomototor response rate is attenuated during tacahycardia. (A) Schematic cartoon of experimental rig for recording tail and heart activity moving gratings (OMR) and salt-pulses via the odor delivery pencil. (B) Example data collected from the heart (pink) and tail (blue) from two OMR trials. In the left trial the fish swims, which terminates the stimulus. In the right trial there is no swim response and the trial ends after 8 seconds. Scale bar indicates 2 seconds. (C) Sample experiment where heart-rate is recorded (pink) while observing the fish’s tail (blue) respond (bottom, light) or not (top, dark) to optomotor trials interspersed with salt pulses (orange). (D) Relationship between heart rate and probability to respond during an 8-second whole-field motion trial. Light gray lines indicate individual fish (n=17). Error bars are SEM. (E) Amongst trials where the fish swims, the relationship between heart rate and time it takes to respond to an 8 second whole-field motion trial amongst OMR trials where the fish swims. Light gray lines indicate individual fish (n=17). Error bars are SEM.

As in earlier studies, we find that while larvae are inactive during interstimulus periods, they often struggle within seconds after the presentation of either cue (Figure S1B-D). These struggles are associated with an increase in heart rate, which otherwise is stable during the interstimulus periods. After a threat, the cardiac response often begins with a brief bradycardia, but is reliably followed by a persistent rise, on average, 0.13 Hz and 0.05 Hz for salt pulses and dark flashes, respectively, before slowly decaying back to baseline over at least a minute (Figure S1B-D). No cardiac modulation was observed when threats failed to induce a struggle (Figure S1E,F). Furthermore, the cardiac dynamics around spontaneous struggles mirror those that occurred during a threat (Figure S1G). Overall, when comparing the relative onsets of all tachycardia events, we find that roughly 95% of these events begin within 10 seconds of a struggle (Figure S1H), suggesting the observed tachycardia states are coupled to high-energy locomotor output rather than the tested stimuli.

### Zebrafish larvae are less responsive to whole-field motion (OMR) during tachycardia

We explored whether these environmentally induced changes in heart rate were associated with changes in the optomotor response (OMR), in which the fish is shown a pattern of moving stripes and tends to respond by moving its tail in the direction of movement. We compared the number of times the fish responded or did not respond to the OMR challenge depending on heart rate (Figures 1B, 1C). We found that both threat-induced and non-stimulated tachycardia was associated with a marked decrease in responsive OMR trials (Figure 1D, Figure S1J). When the larvae did respond during tachycardia, their accuracy was unaffected (Figure S1I), but their latency to respond was longer, suggesting that the animal is raising its threshold to swim (Figure 1E). We found the same inverse correlation between heart rate and responsiveness to dark flashes, indicating a non-specific change in behavioral state (Figure S1K).

Of course, the changes in OMR and tachycardia, although correlated, may be independent responses to the challenges or to the struggle itself. We thus sought to disentangle the struggles from tachycardia events by directly manipulating the sympathetic nervous system, pharmacologically and optogenetically.

### Sympathetic manipulation simultaneously impacts cardiac and behavioral states

We first asked whether the sympathetic nervous system is indeed activated during the threat-evoked tachycardia state. To that end, we performed calcium imaging in transgenic fish line, *tg(Th1:gal4; UAS:GCaMP6f)*, that express a calcium indicator in noradrenergic (and dopaminergic) neurons^31^ (Figures 2A, 2B). These experiments revealed that neuronal dynamics across both bilateral pairs of sympathetic ganglia strongly reflect the heart rate (Figures 2C–2E). Nearly every neuron imaged within these ganglia shows a high correlation (>0.5) with heart rate (Figures 2D, 2E).

**Figure 2.**
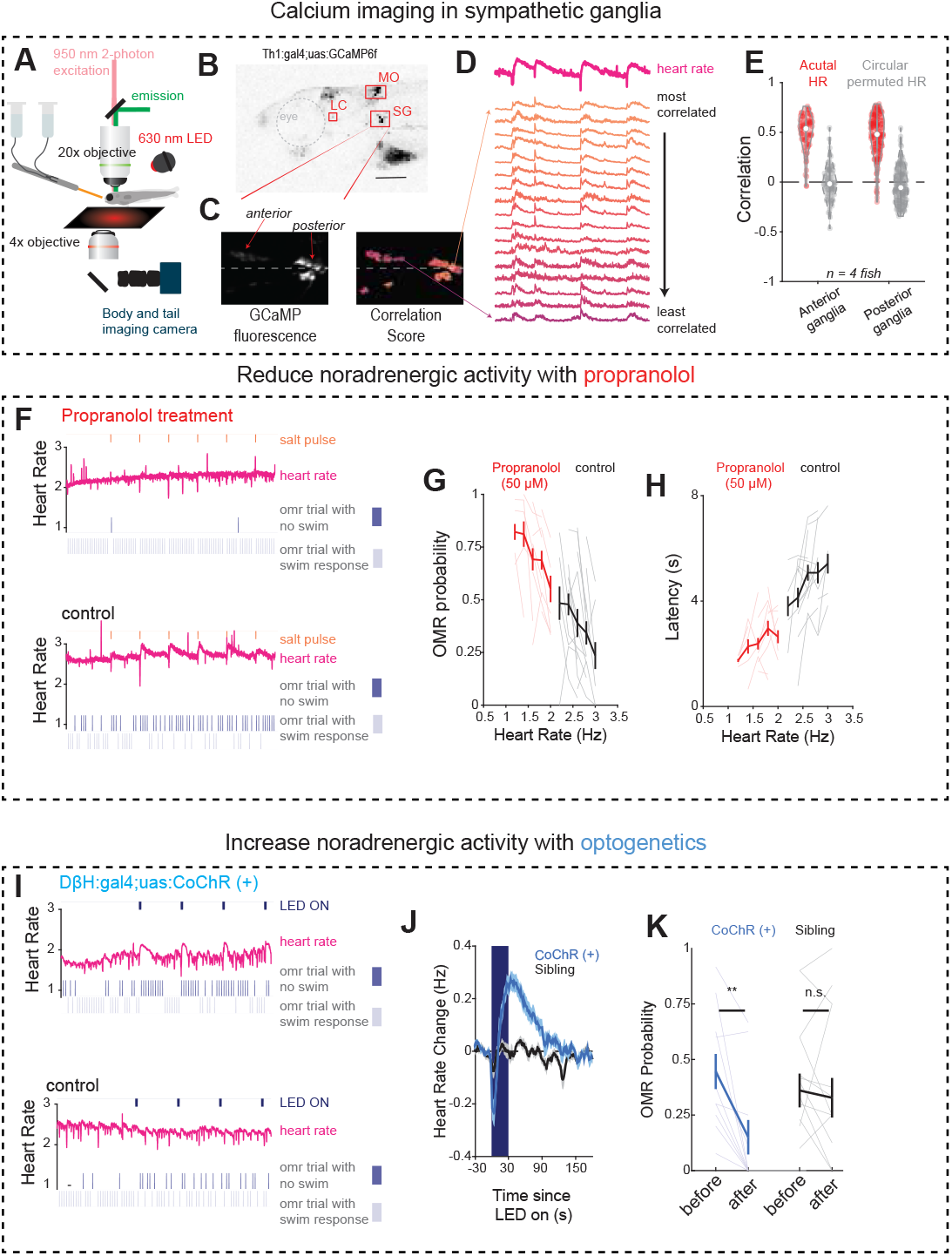
Noradrenergic signalling encodes and regulates cardio-behavioral state. (A) Schematic cartoon of experimental rig for imaging the brain, tail, and heart, while also presenting dark flashes (via the red LED) and salt-pulses via the odor delivery pencil. (B) Projection of confocal stack of *th1:gal4;uas:GCaMP6f fish* imaged from the side. Eye indicated by dashed circle. scale bar (150 *µ*m) (C) Dorsal view of a 2-photon image of the sympathetic ganglia anatomy (left) and correlation score with heart rate for each cell’s calcium signal overlaid on top (right) and indicated by color. scale bar (50 *µ*m) (D) Heart rate and calcium traces from (C) sorted by correlation score. (E) Correlation of all sympathetic neurons in the anterior (left) and posterior (right) ganglias to recorded heart rate (red) and circularly permuted heart rate signal (gray). (F) Sample experiment recording heart rate and OMR performance (see Figure 1C) in a fish treated with 50 *µ*M propranolol (top) and a sibling control (Bottom). (G) Relationship between heart rate and probability to perform OMR by fish treated with 50 *µ*M propranolol (n = 13) and sibling controls (n = 15). Light lines indicate individual fish. Error bars are SEM. (H) Relationship between heart rate and latency to first swim during OMR trial by fish treated with 50 *µ*M propranolol (n = 13) and sibling controls (n = 15). Light gray lines indicate individual fish. Error bars are SEM. (I) Sample experiment recording heart rate and OMR performance while optogenetically activating noradrenergic neurons (blue bars at top) in a *DβH:gal4;uas:CoChR-eGFP* fish (top) and an *eGFP*(−) sibling control fish (bottom). (J) Heart rate change elicited by blue light exposure to *DβH:gal4;uas:CoChR-eGFP* fish (blue, p=0.00011, paired t-test, n = 10) and an *eGFP*(−) sibling control fish (gray, p = 0.5837, paired t=test n = 11). Shaded indicates SEM. (K) Change in optomotor response rate between the 90 seconds before blue light exposure and 90 seconds after blue light exposure for *DβH:gal4;uas:CoChR-eGFP* fish (blue, p =0.002, paired two-sided rank test, n = 10) and *eGFP*(−) sibling control fish (gray, p = 0.4667, paired two-sided rank test, n = 11). Lighter lines indicate individual fish. Error bars are SEM.

Blockade of the effects of norepinephrine on the heart using propranolol (Figure 2F) led to a 55% reduction of the average baseline heart rate across fish (Figure S2A) that was accompanied by a 40% increase in baseline optomotor response probability Figure S2B). Stressors continued to generate struggles, but the tachycardia that followed was blunted by 40% after propranolol (Figure S2C) along with an increase in optomotor responsiveness (Figure 2G). Interestingly, the residual variation in heart rate was still predictive of optomotor response probability and latency (Figures 2G, 2H). In fact, treated fish appear to occupy the same linear function that connects heart rate to these parameters in controls.

To separate sympathetic activation from the naturally co-occurring struggles, we optogenetically excited the sympathetic ganglia of transgenic fish expressing CoChR in cells that produce dopamine-β-hydroxylase, an enzyme necessary for converting dopamine into norepinephrine (Figure 2I). At baseline, these fish behave similarly to controls (Figure S2D,E). We observed that brief illumination with blue light did not cause the fish to struggle (Figure S2F). It did, however, induce a strong and long-lasting (90 s) tachycardia (Figure 2J). Wild-type controls showed no behavioral or cardiac responses to the blue light. Furthermore, during this optogenetically induced tachycardia, the likelihood of the fish to respond to OMR is significantly reduced by 65% (Figure 2K). This is not true for control fish during the minute that follows blue light exposure. Interestingly, across control and stimulated fish, heart rate and the optomotor response occupy the same overall relationship – further supporting a tight relationship between the two (Figure S2G,H). These data indicate that sympathetic stimulation is, not surprisingly, likely to underlie the tachycardia induced by threat. In this case we find that the changes in heart rate are associated with changes in OMR responsiveness, regardless of whether or not accompanied by struggle. Yet, it is still possible that sympathetic activation drives heart rate and behavioral changes independently. Hence, we sought to directly change heart rate in order to see whether that sufficed to alter OMR responsiveness.

### Slowing heart rate by blocking cardiac pacemaker activity increases optomotor response rate

We first pharmacologically inhibited cardiac pacemaker activity using the HCN4 blocker ivabradine (Figures 3A–3C)^32–34^. We found that treatment with ivabradine dramatically reduced the heart rate (−46% Figure S3A), and increased the optomotor response (91% Figure S3B) (Figure 3B) and decreased OMR latency (Figure 3C). Furthermore, we found that the HCN4 blockade had minimal effects on the change in heart rate after a threat (Figure S3C), indicating that the underlying mechanisms that drive tachycardia are still intact. We replicated these findings by testing a second drug, ZD7288 (Figure S3D-H) that also inhibits HCN4. Together, these results strongly suggest that it is the heart rate, per se, that influences OMR responsiveness, even in the absence of struggle or sympathetic activation.

**Figure 3.**
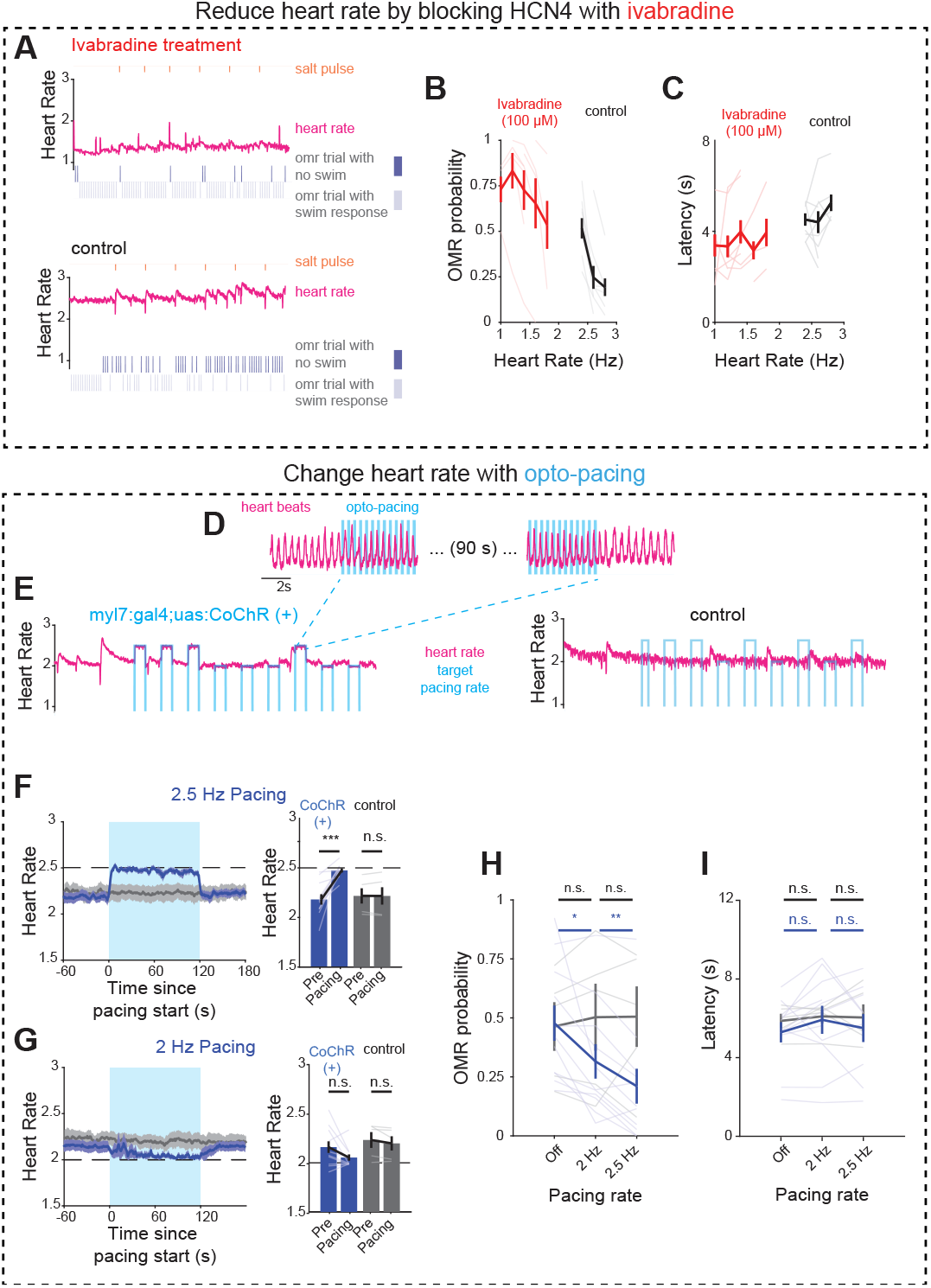
Direct manipulation of cardiodynamics impacts the optomotor response. (A) Sample experiment recording heart rate and OMR performance (see Figure 1C) in a fish treated with 100 *µ*M of the HCN4 inhibitor ivabradine (top) and a sibling control (bottom). (B) Relationship between heart rate and probability to perform OMR by fish treated with 100 *µ*M ivabradine (n = 7) and sibling controls (n = 6). Lighter lines indicate individual fish. Error bars are SEM. (C) Relationship between heart rate and latency to first swim during OMR trial by fish treated with 50 *µ*M propranolol (n = 7) and sibling controls (n = 6). Lighter lines indicate individual fish. Error bars are SEM. (C) Zoom in of pacing period in (E) showing individual heart beats 5 seconds before and during pacing (blue bars indicating times where blue light is on) and 5 seconds after pacing stops. (D) Sample experiment recording heart rate while optogenetically pacing the heart to different target rates (2 or 2.5 Hz, blue line) in a *myl7:CoChR-tdTomato* fish (left) and an *tdTomato*(−) sibling control fish (right). (E) Left : average time course of heart rate centered on onset of 2.5 Hz pacing for *myl7:CoChR-tdTomato* fish (blue, p = 0.000374, paired t-test) and *tdTomato*(−) sibling control fish (gray, p = 0.9770, paired t-test). Shaded area indicates SEM. Right: average heart rate before and during pacing for the same groups. (F) Left : average time course of heart rate centered on onset of 2 Hz pacing for *myl7:CoChR-tdTomato* fish (blue, p = 0.0726, paired t-test) and *tdTomato*(−) sibling control fish (gray, p = 0.2102, paired t-test). Shaded area indicates SEM. Right: average heart rate before and during pacing for the same groups. (G) Relationship between optical pacing and probability to perform OMR in *myl7:CoChR-tdTomato* fish (blue, n = 10, p = 0.0182 and p = 0.0030, paired t-test) and tdTomato(−) sibling controls (gray, n = 5, p = 0.3712 and p = 0.9762, paired t-test). Lighter lines indicate individual fish. Error bars are SEM. (H) Relationship between optical pacing and latency to first swim during an OMR trial in *myl7:CoChR-tdTomato* fish (blue, n = 10, p = 0.1465 and p = 0.5562, paired t-test) and *tdTomato*(−) sibling controls (gray, n = 5, p = 0.3799 and p = 0.9204, paired t-test). Lighter lines indicate individual fish. Error bars are SEM.

### Optogenetically paced tachycardia blunts the optomotor response

To directly control heart rate, we adopted a technique, optopacing, that has been demonstrated in zebrafish^35^ and mice^8^, by expressing an optically gated cation channels (CoChR) in cardiomyocytes (myl7) and using timed illumination to pace the heart at set frequencies (Figures 3D, 3E). We selected two target heart rates, 2 Hz and 2.5 Hz, the lower rate corresponding to a value near the baseline heart rate, and the higher value closer to the peak of stimulus-evoked tachycardia (Figures 3F, 3G). We found that pacing at the baseline rate itself diminished the OMR response, but not latency (Figures 3H, 3I). We presume that this diminution may be due to the effect of changing the pattern of cardiac contraction and changes in cardiac output, since the induced heart beat may originate from outside of the normal pacemaker region. Even so, as shown in Figure 3H, increasing the rate of stimulation progressively reduced the OMR response, as predicted. In control larvae exposed to the same light stimulation, OMR responses are unaffected (Figures 3H, 3I)). Hence, under each situation – external stimuli, sympathetic stimulation, pharmacological treatment, and opto-pacing – OMR decreases as heart rate increases. This indicates that some attribute of heart rate is sensed by the nervous system.

### Neuronal dynamics reflect heart rate changes

In order to begin to understand how and where information about heart rate is captured and transmitted to CNS centers responsible for behavioral modulation, we explored neuronal activation in the nodose and epibrachial ganglia (vagal) and across the CNS (Figure 4A). We first recorded calcium activity from vagal sensory neurons of *tg(vGlut2a:gal4;UAS:GCaMP6s)*^36^ fish while presenting threats and recording cardiac and behavioral responses (Figure 4B). The nodose ganglia receive afferents not only from the heart but from many visceral organs. From these data, as found recently^37^, we uncovered many primary sensory neurons with dynamics that correlate with heart rate (Figure 4B, S4A). These were widely distributed across the different vagal ganglia (Figure 4C). Since the epibranchial ganglia primarily innervate the gills, and the nodose receives most of its inputs from other visceral tissues, some fraction of heart rate related activity in the nodose likely originates from sources other than the heart.

**Figure 4.**
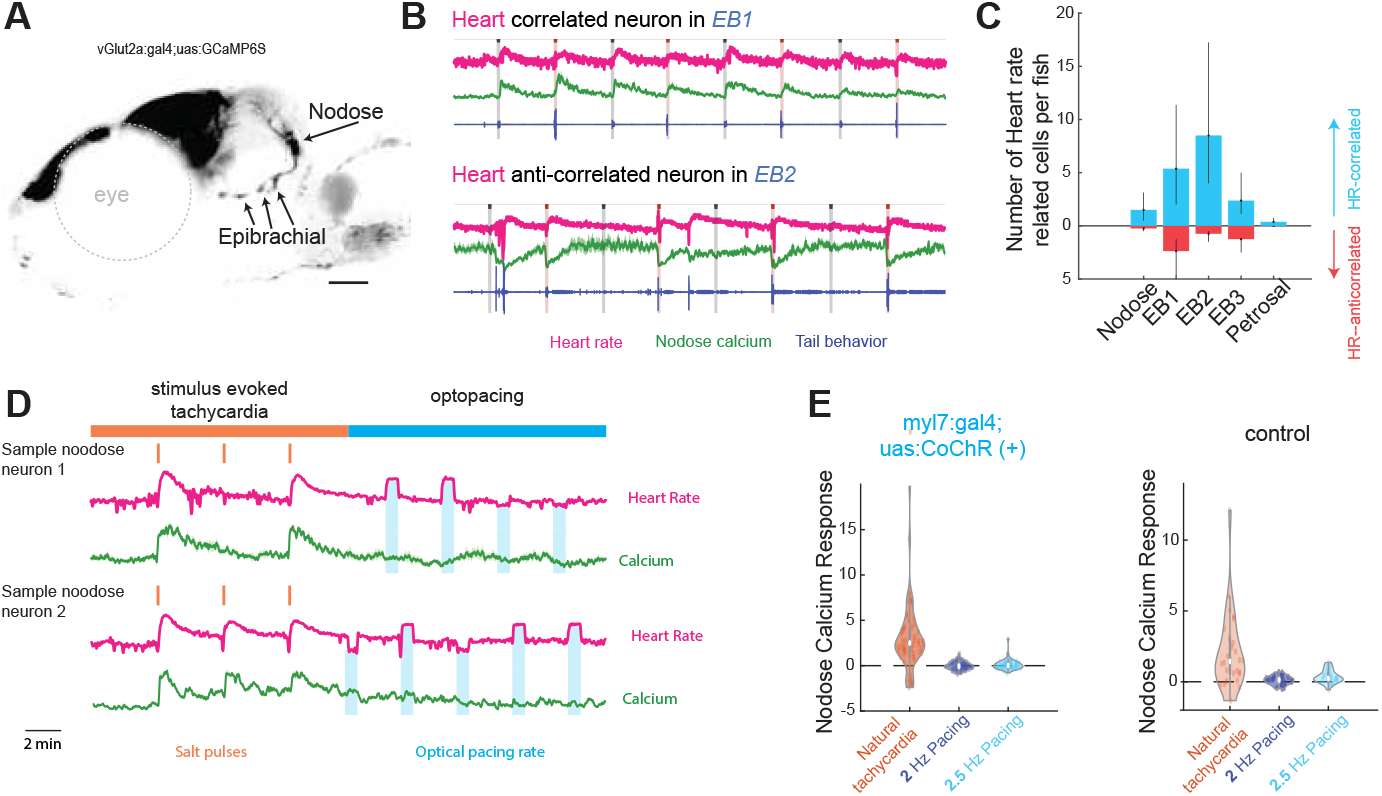
Activity in vagal sensory ganglia reflect stimulus-evoked cardiac state, but not optogenetic cardiac pacing. (A) Projection of confocal stack of *vGlut2a:gal4;uas:GCaMP6s* fish imaged from the side. Eye indicated by dashed circle. scale bar (150 *µ*m). (B) Sample heart rate, behavior and calcium activity traces recorded from heart rate correlated (top) and anticorrelated (bottom) cells from the epibrachial ganglia. (C) Median number of heart rate correlated (r > 0.3, blue) and anti-correlated (r < −0.3, red) cells found in each ganglia (Nodose, 3 epibrachial, and petroal) across 8 fish. Error bars indicate 95% confidence intervals (1000 bootstraps). (D) Sample recordings of heart rate and calcium from an experiment combining optopacing and calcium imaging in the nodose showing cells that are correlated with heart rate during “natural tachycardia” epoch and the responses of those cells during the 15 minute optopacing period of 60 seconds of pacing separated by 120 seconds of rest. (D) Violin plots of the distribution of the activity of “natural heart-rate” correlated neurons following presentation of salt pulse (orange), 2 Hz pacing (dark blue), and 2.5 Hz pacing (light blue) for *myl7:CoChR-tdTomato* fish (left, n = 4 fish) and *tdTomato*(−) sibling controls (right, n = 4 fish).

To explore how much, if any, heart rate information was coming directly from the heart, we used the optopacing technique to change heart rate while simultaneously monitoring vagal sensory activity. Using this approach we failed to identify vagal sensory activity that responded to optopacing (Figures 4D, 4E). This suggested that the nodose may not be the path by which afferent information about heart rate is transmitted to the CNS.

We next imaged activity of glutamatergic neurons across the brain using 2-photon microscopy which we registered to the Z-brain atlas^38^ in order to map activity to specific brain regions (Figures S4, 5A). We observed cells in particular regions whose activity were either strongly correlated or strongly anti-correlated with heart rate (Figure 5B). The areas with activity anti-correlated with heart rate included the pallium, thalamus, habenula, and inferior olive among many others (Figures 5C–5E). Most prominent among these anti-correlated regions is a thalamic structure labeled “Mesencephalic Vglut cluster 1” that has previously been compared to the mammalian Edinger-Westphal nucleus and has been connected to regulating response latencies and arousal states^29^.

**Figure 5.**
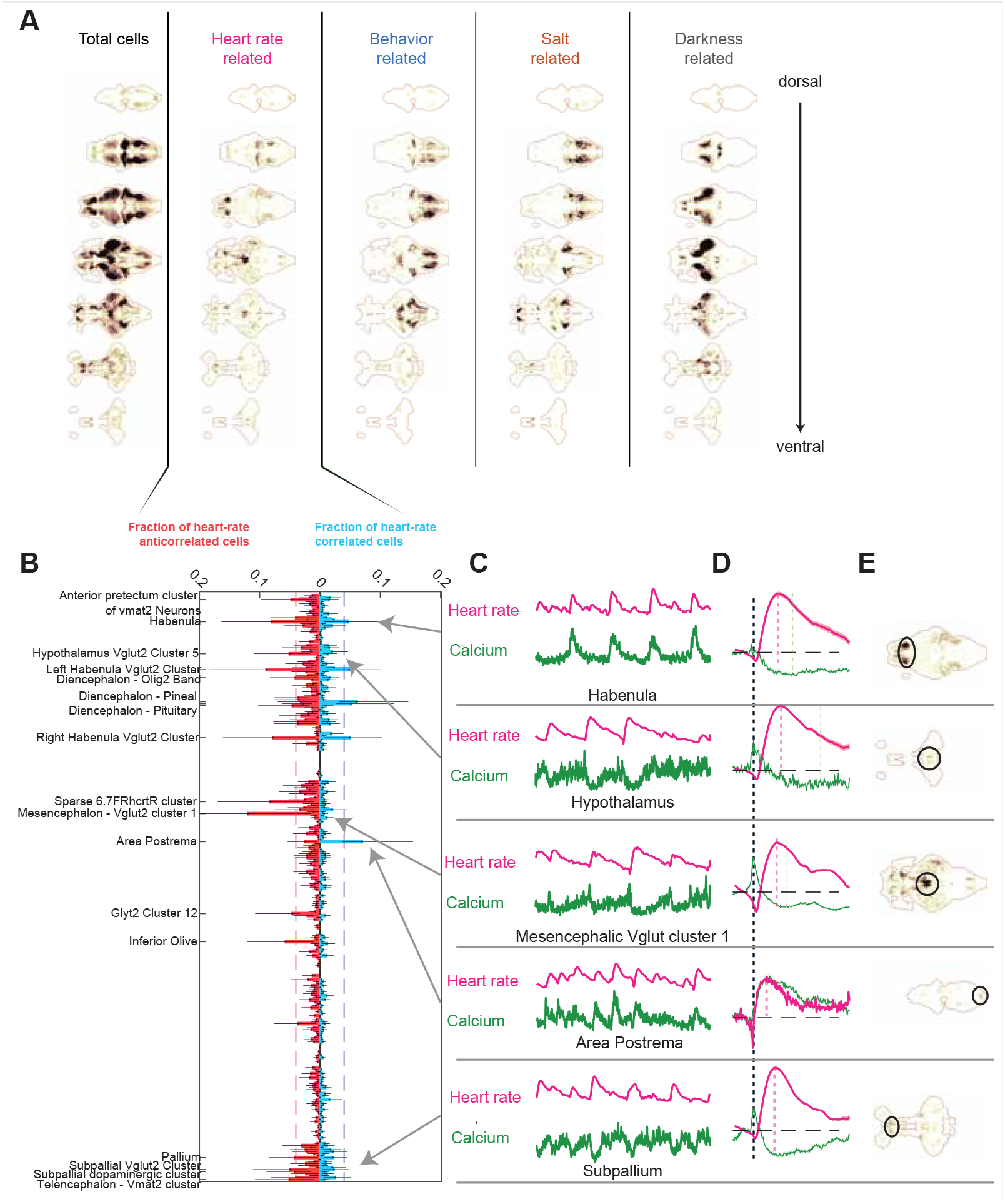
Cardiac state is encoded in neural dynamics across distributed brain regions. (A) Brain wide maps of all recorded neurons aross *vglut2a:gal4;uas:GCaMP6S* fish registered to the Z-Brain atlas, heart-rate correlated neurons, as well as behavior, salt, and dark flash correlated neurons. (B) Fraction of significantly heart-rate correlated (blue,right) and anti-correlated (red,left) cells in every Z-brain demarcated mask. Labeled masks are those containing the highest fraction of heart-rate encoding cells (>5% of cells is either correlated (r > 0.3) or anticorrelated (r < −0.3). Error bars indicate 95% condfidence interval. (C) Sample heart rate and calcium activity traces from representative cells in the area postrema, habenula, hypothalamus, mesencephalic vGlut cluster 1, and subpallium. (D) Struggle-triggered average of heart rate and calcium dynamics from heart-rate correlated cells in the areas indicated in (C). (E) Anatomical location in (A) of the regions depicted in C-D.

Of all the regions we imaged, we found the area postrema to have the highest frequency of neurons positively correlated with heart rate (Figure 5B). The area postrema is of interest because it is a circumventricular organ lacking a blood-brain barrier and therefore particularly sensitive to circulating peptides, including those regulating hemodynamics such as angiotensin, and believed to include circuits critical to regulation of blood pressure and heart rate^39–41^. We also found that, unlike neurons of the nodose, activity in the area postrema increased dramatically during tachycardia induced by direct cardiac pacing (Figures 6A, 6B). Mirroring its effects on the optomotor response, pacing at both 2 and 2.5 Hz increased area postrema activity (Figure 6C). These data suggest one possible route by which changes in heart rate may influence brain activity and ultimately modulate the behavior of the animal.

**Figure 6.**
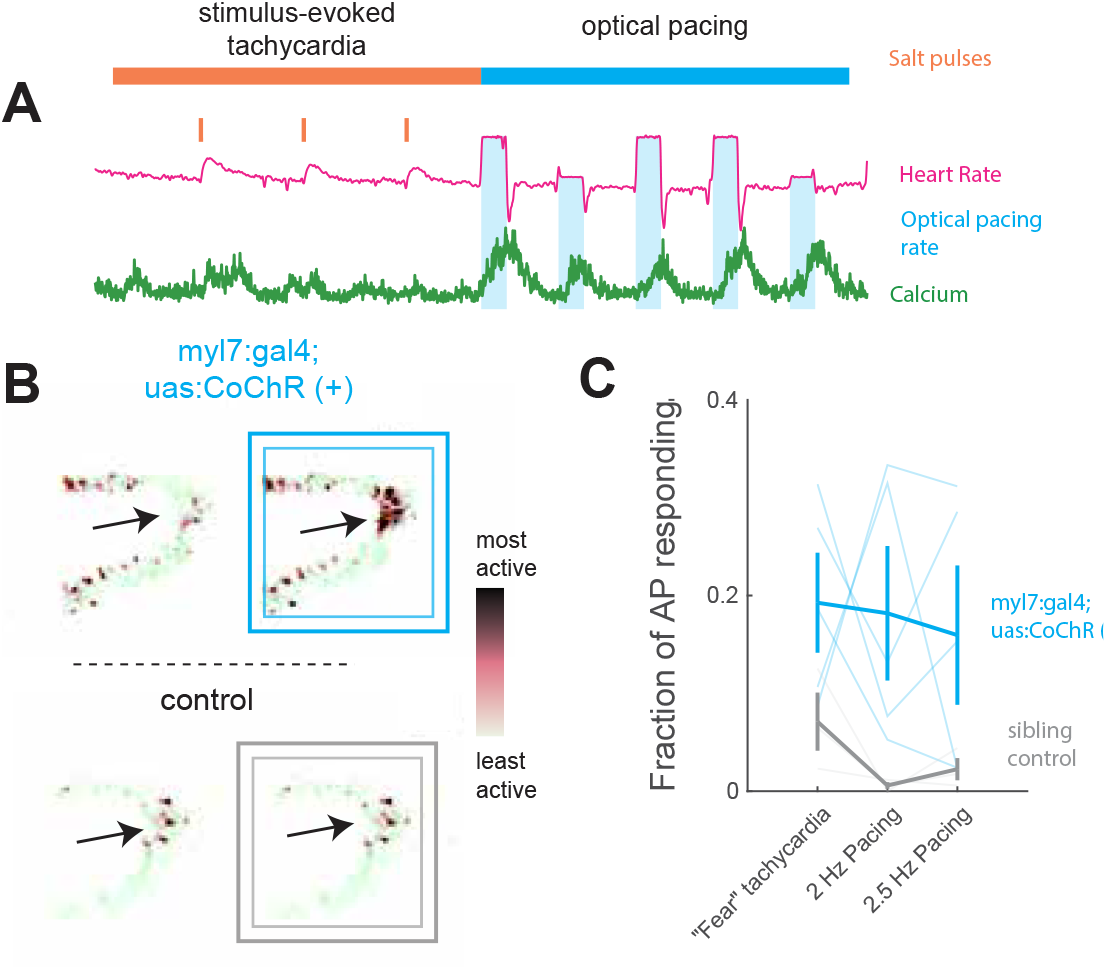
Excitatory activity in the area postrema is increased by manipulating heart rhythm. (A) Sample recording of heart rate and calcium from an experiment combining optopacing and calcium imaging in the nodose showing cells that are correlated with heart rate during “natural tachycardia” epoch (orange) consisting of 15 minutes of 10 second salt pulses separated but not during the 15 minute optopacing period of 60 seconds of pacing separated by 120 seconds of rest (blue). (B) Raw fluorescence of area postrema during baseline and pacing (squares) for *CoChR-tdTomato* (blue) and control fish (gray). Activity increases from light to dark. (C) Fraction of area postrema that is active during pacing for *CoChR-tdTomato* (blue) and control fish (gray).

## DISCUSSION

We have found here a correlation between heart rate and a simple sensori-motor task, the optomotor response, in larval zebrafish. Regardless of the drive to heart rate change, whether environmental threat, pharmacological manipulation, or direct optogenetic pacing of the heart – the faster the heart rate the lower the optomotor response rate. We find neurons that are activated when the heart goes more quickly (tachycardia) and others activated when it goes more slowly (bradycardia). These include afferent sensory cells in the nodose ganglion and in specific neural centers of the CNS. This suggests that heart rate is assessed and used to modulate particular neural circuits and behaviors.

Classically, interoception and motor control of the cardiovascular system is viewed through the lens of maintenance of internal homeostasis. Two parameters are assessed, blood pressure by baroreceptors and oxygen partial pressure by chemoreceptors, and heart rate and cardiac contractility adjusted in the face of challenge primarily by autonomic output^2,4,42^. In this scenario, heart rate is viewed as a component of the motor limb, helping to adjust cardiac output to changing demands of the body. Here we find that heart rate may be assessed as part of the afferent limb. This finding adds to evidence that changes in heart rate affect broader brain state^43,44^. A century ago, James and Lange proposed that changes in internal functions, including heart rate, can drive emotions, such as fear. Recent work, directly modulating heart rate in mice, would support this notion^8,45^. By uncovering this phenomenon in a developing teleost we open the door to unraveling new mechanistic insights more readily available in this fully optically accessible model.

While it is clear that specific neurons encode tachycardia or bradycardia, we do not know the cellular mechanisms that enable heart rate reception, and whether it acts via direct or indirect pathways. We do not believe that all, if any, heart rate-responsive neurons innervate the heart because we find heart rate responsiveness in the branchial ganglia, where neurons have been only reported to innervate the gill arches. It is conceivable, for example, that vascular pulsatility through endothelial signaling^46^ or fluid dynamics^47^ is the driver. This might also be the case in the responsive CNS neurons, particularly in those populating the area postrema, which lacks a blood-brain barrier and is in direct contact with the vasculature^48^. No matter which mechanism and cell type is used, individual beats need to be integrated with extended time constants in order to transform temporally spaced binary events into an analog value that encodes frequency. This requirement constrains physiological mechanisms that must involve time constants exceeding the capacity of individual neurons and may be found either in distributed circuits^49,50^ or in specialized non-neuronal cell types such as radial astrocytes^27,51^.

Here, we explored a quantitative sensorimotor response, the optomotor response (OMR), whereby a larval fish makes a decision to move in the direction of whole-field motion. The circuit underlying OMR is well-defined at the cellular level, from its sensory inputs in the retina to sites of decision-making in the anterior hindbrain and motor control among the ventromedially located spinal projection neurons. This system allowed us, in an unanesthetized animal, to quantify the relationship between heart rate and a specific reflex response and to begin to define the neural circuitry responsible for integrating heart rate with sensorimotor control. The OMR depends on temporal integration of motion by cells in the anterior hindbrain^24^ to determine whether an animal turns in response to the visual input, and the latency before it does so. We find here that tachycardia reduces the response rate and increases the latency, but does not affect the accuracy of turning (i.e., the animal turns in the “correct” direction), suggesting the rates of this integration may be influenced by cardiac state. We do not yet know which of the heart rate-responsive neurons interact with the hindbrain circuits responsible for the OMR. However, some of these regions are known to be involved in regulating behavioral states. For example, we find populations of heart rate anti-correlated neurons across the preoptic area of the hypothalamus and Edinger-Westphal nucleus. These are both regions that are important for managing arousal^29,52^ and energy homeostasis^53,54^. The habenula, through its projections to the interpeduncular nucleus^55^ and raphe^56,57^, is a critical hub for managing behavioral selection in fish and mammals^58–60^. How these various neural populations interact is unknown, but will become clear as comprehensive wiring diagrams of the zebrafish brain emerge from connectomics efforts^61–64^.

Why might animals have evolved circuitry to couple cardiac changes to behavior? One possibility is that heart-rate may act as a proxy for energy debt. The tachycardia events we have studied, with the exception of when we apply certain optogenetic perturbations, are minutes-long periods that follow vigorous struggle attempts. This period is likely accompanied by an energy deficit. If the heart rate reflects the collective homeostatic mechanisms that are bringing the animal back to its resting state, then it would be a valuable variable to monitor to halt unnecessary expenditure. In such a scenario, emotions may represent an evolutionary offshoot of an ancestral condition where animals used behavioral modulation to preserve physiological homeostasis.

## METHODS

### Data acquisition

#### Animal husbandry

All live animal experiments were performed according to protocols approved by Harvard University Institutional Animal Care and Use Committee (IACUC). For all transgenic fish used, crosses were made between adults that were heterozygous carriers of a mitfa (+/-) mutation that generates the nacre phenotype^65^. For all experiments, mitfa (−/-) null mutations were screened for and utilized. Larvae were housed in an incubator at 28°C with a 14/10 hr day/night cycle until 5 days post fertilization (dpf) in filtered facility water. The water in dishes containing embryos was exchanged daily. After 5 dpf, larvae were fed fresh paramecia daily. All experiments were performed on larvae between 5 and 7 dpf. A list of transgenic lines used is provided in the resource table.

#### Transgenic line generation

The *tg(myl7:CoChR-tdTomato)* line was generated using the tol2 transposase (cite) method. Briefly, gateway cloning was used to combine a p5e vector containing the 3kb myl7 promoter, a pMe containing *CoChR-tdTomato*^66^ and a p3e containing a poly-A tail. This plasmid was injected into the single-cell staged embryos of *tg(vGlut:gal4;uas:GCaMP6S)* parents. Embryos were screened for *tdTomato* expression and propagated.

#### Heart rate and behavior acquisition

For all behavior and imaging experiments, larvae were embedded in 2% low melting point agarose on 50 mm petri dishes. Following embedding, the fish and agarose were submerged in filtered facility water and allowed to further stabilize for at least 2 hours. Approximately 30 minutes before the experiment began, the agarose surrounding the tail, as well as the rostral tip of the fish were removed to allow for trackable tail movements and chemical stimulus presentation, respectively. Salt stimuli (50 mM NaCl) were presented to the fish using a previously described gravity-driven perfusion system^28^. Visual dark flashes were presented through a 625 nm LED (Thorlabs M625L4) placed above the fish. All stimuli were presented for 10 seconds, with interstimulus intervals of either 300 s or 60 s.

Tail tracking and heart rate recordings were acquired through a single FLIR Blackfy camera placed below the fish at 100 Hz via infrared illumination (850 nm ThorLabs LED M850L3) presented from above the fish. In the light path between the IR LED and the camera, we placed a 4X objective (AMScope PF4X-INF) to provide enough magnification to visualize the heart. Before the experiment begins, the center location of the heart is manually selected, and a 20×20 pixel image surrounding the heart is saved for further processing every 30 ms. For tail tracking, a tail skeleton consisting of 10 points is automatically detected and saved using a previously described algorithm^67^ at every time point (100 Hz) during the experiment. Stimulus control, as well as heart and behavior acquisition, were performed using a single custom-written LABVIEW program.

For combined heart-rate and optomotor response experiments, a second preparation was built, with the objective and camera placed above the fish and infrared illumination placed below. This allowed for a projector to be placed below the fish in order to present visual stimuli.

To induce fear states, we used the chemical delivery apparatus described above. The square gratings moved to the left or right for up to 8 seconds every 20 seconds. To prevent struggles being induced by perceived futility, we stopped the moving gratings after each turn event. To induce heart rate dynamics comparable to the ones generated in the other assays, we presented 10-second pulses of 50 mM salt to the larvae every 6 minutes.

#### Whole field motion stimulation

To probe the larvae’s visual sensitivity during tachycardia, we presented black and white gratings with a period of 7 mm to the scene directly below the fish using custom-written LabVIEW software. To measure heart rate in these experiments, we placed the recording camera and 4X objective above the fish. Visual gratings moved at 1.5 cm/s either to the left or right every 20 seconds for up to 8 seconds. To prevent futility-induced passivity^27^ grating motion ceased immediately after the fish attempted a turn as detected by the online tail-tracking software

#### Pharmacology

All working solutions were made fresh in filtered fish water the same day of the experiment. Larvae were first incubated in the drug for at least 2 hours and then embedded. The embedded larvae were then immersed in the same concentration of drug. Sibling control fish went through the same procedure, but without any drug added to the water.

#### Calcium Imaging

For all functional imaging experiments, we used a custom-built two-photon microscope. Excitation was provided by a femto-second pulsed Ti:Sapphire laser (MaiTai, Spectra-Physics) set to 950 nm. The sample at power was <10mW. Frames were acquired at 0.7-1.1 Hz. Microscope control and image acquisition were performed by a single custom-written LABVIEW program. For nodose imaging experiments, larvae were embedded on either their right or left sides. For all other experiments, larvae were embedded dorsal side up.

#### Optogenetic cardiac pacing experiments

To perform optical pacing concurrent with testing their optomotor response, a slit was cut into the side of a 10 mL petri dish to allow the cannula of a ThorLabs fiber optic cable (M98L01) to be fixed flush with the floor near the center. The wall was repaired with epoxy. Larvae were embedded such that the output of the fiber-optic cable was even with the posterior side of the heart. A power was identified that did not affect control fish, and still led to cardiac contractions in transgenic fish positive for *myl7:CoChR*. During pacing sessions where the target frequency was 2 Hz, the LED was on for 200 ms and off for 300 ms. For 2.5 Hz the LED was on and off for 200 ms.

During imaging experiments, we needed to avoid illuminating the fish and recording light from the photomultiplier tube at the same time to prevent photodamage. Thus, we used a gated PMT (Hamamatsu H11526-01-NN) for collecting emitted photons, and interleaved recording and stimulation during the galvometer scanning, such that during the forward scan, the PMT was on and light source off, and during the backward scan, the light source was on and PMT unpowered. To enable rapid switching between on and off states, we used a blue laser (Coherent OBIS 1178767). This was fed directly to the heart of the fish via a 400 *µ*m fiber optic cannula (CFMLC14L20) - controlled by a manual micromanipulator (Siskiyou SD-130). The pacing stimulation was otherwise the same as for the behavior.

### Data analysis

#### Extraction of heart rate from saved images

To extract heart rate from the 20×20 pixel movies saved during all experiments, we used a custom pipeline written in MATLAB. We first performed PCA across time and then examined the first 7 principal components to identify the components with the strongest frequency power in the 0.5-5 Hz band. This signal was assumed to be related to heart rate. We then extracted the time between peaks in this signal using MATLAB’s findpeaks function and manually selecting a ‘MinPeakProminence’ value that sufficiently captured peaks after visual inspection. Peaks detected within 200 ms of the preceding peak were removed. The instantaneous heart rate was then defined as the inverse of the time between the detected peaks. All fish with a median basline less than 1 Hz were removed from analysis.

#### Image segmentation and registration

Neural signals from within the brain were extracted by first segmenting every imaged plane into 2-dimensional voxels. This was done by first correcting for motion in XY across time^68^ and then performing segmentation based upon local correlations of pixel fluorescence across time following a previously described protocol^28,69^. For the sympathetic nervous system - which has relatively few cells - neuronal cell bodies were manually segmented. Registration of functional stacks to a single reference volume was done using the BigWarp plugin in FIJI^70^. Landmarks that connected functionally imaged anatomy to the reference anatomy were selected individually for each experiment.

#### Identification of heart-rate related cells

To classify neuronal responses we first computed Δ*F/F* values, assuming the baseline to be the median fluorescent values. We then specifically selected cells that were “active” above noise by comparing the variance of the frame-by-frame differential of the trace to the overall trace. Cells where this value was greater than 1.2 were deemed to be inactive or noisy, and removed from analysis. We then generated regressors from the heart rate and tail behavior of the animal, as well as the stimuli presented to the animals. Each parameter was convolved with a kernel representing the onset and offset dynamics of GCaMP6S to make these regressors. These regressors were cross-correlated with each neuron’s calcium trace. Neurons were defined to be most correlated to a given stimulus when that stimulus generated its greatest correlation and the peak correlation coefficient was greater than 0.3.

## Data and Code Availability

Data and code will be made available upon publication.

## Acknowledgements

NIH grant U19NS104653 and 1R01NS124017 to F.E. and M.C.F.; Howard Hughes Medical Institute to M.B.A.; and Simons Foundation SCGB 542943SPI to F.E. and M.B.A. We thank the Harvard Center for Biological Imaging (RRID:SCR_018673) for infrastructure and support. We thank the Janelia Visiting Scientist Program for support. We thank the Harvard Center for Biological Imaging (RRID:SCR_018673) for infrastructure and support. We thank all members of the Fishman, Engert, and Ahrens laboratories for feedback over the course of this project.

## Author Contributions

Conceptualization, K.J.H., M.C.F., and F.E.; methodology, K.J.H. and A.Z; investigation, K.J.H; visualization, K.J.H.; writing – original draft, K.J.H, M.C.F.; writing – review & editing, K.J.H., F.E., M.C.F.; funding acquisition, M.B.A., F.E., and M.C.F.; project administration, F.E. and M.C.F.; supervision, M.B.A., F.E., and M.C.F.

## Competing Interests

The authors declare no competing interests.

## Supplemental Figures

**Figure S1.**
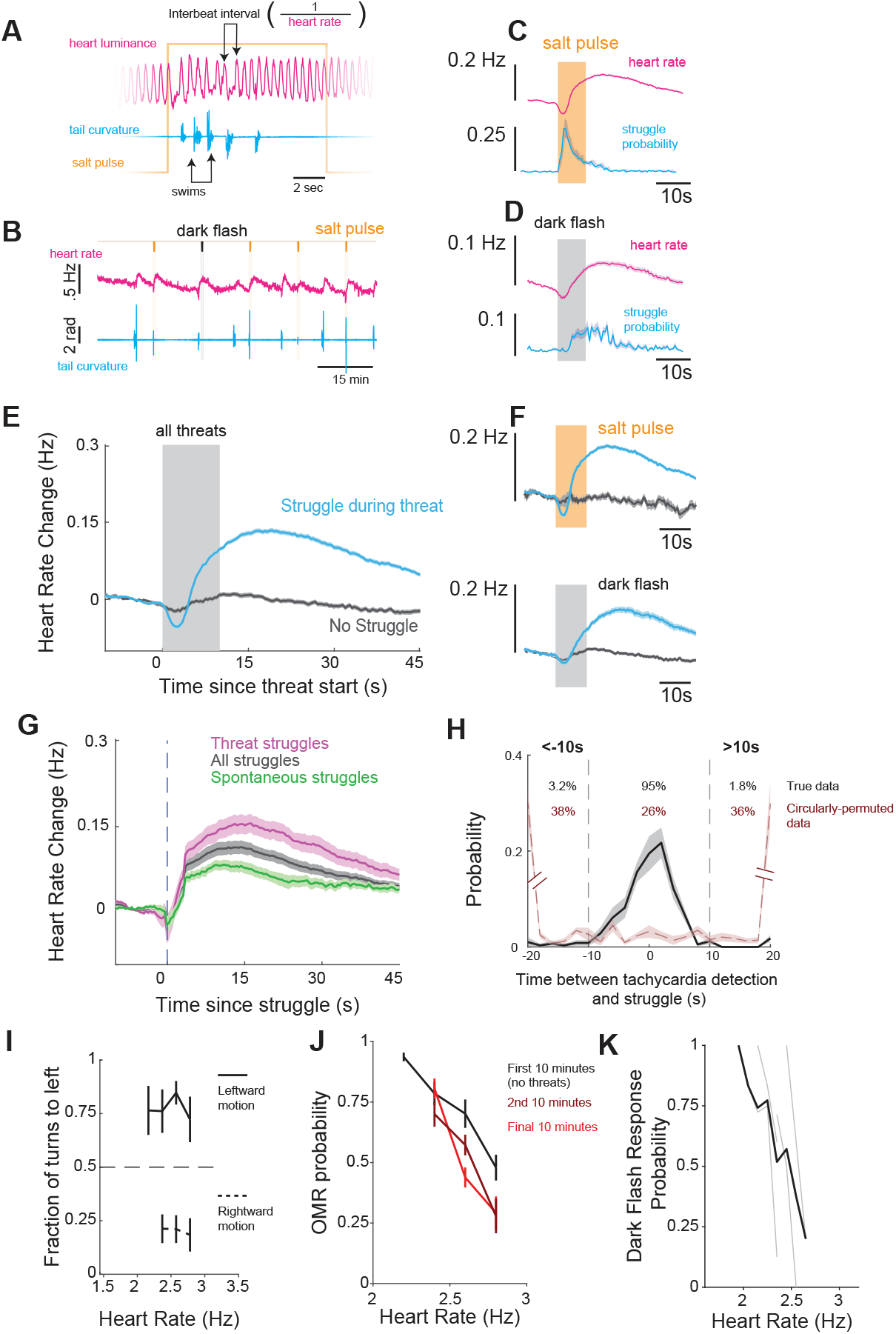
Supporting Material for Figure 1. (A) Sample data from 12 seconds illustrating indvidual heart beats (pink) and tail events (blue) during a threat (salt, orange). (B) Sample experiment showing different threats (darkflash, black, and salt pulse orange), heart rate (pink) and tail (blue) over 30 minutes. (C) Stimulus triggered heart rate (pink) and tail response probability (blue) following a salt pulse. (D) Stimulus-triggered heart rate and tail response probability (blue) following a dark flash. (E) Stimulus triggered heart rate following threats (either dark flash or salt pulse) where the fish struggled during the threat (blue) and those where it did not (gray). (F) Data in (E) split for salt pulses (top) and dark flashes (bottom). (G) Struggle-triggered heart rate average for all struggles (gray), those that ocurred during threats (pink) and those that occurred spontaneously (green). (H) Histogram of the onset of tachycardia events relative to the time of the nearest struggle. Events separated greater than or less than 20 or −20 seconds are accumulated in the largest and smallest bins. (I) Probability of the first turn being toward the left as a function of heart rate (x-axis) and whether or not the motion stimulus is to the left (top, solid) or right (bottom, dashed) (J) Probability of responding to OMR trial versus heart rate for data separated into 3 epochs - first 10 minutes (black), during which no salt-pulses are delivered, and the second (brown) and final (red) 10 minutes, during which salt pulses are delivered every 6 minutes. (K) Probability of a response to a 10 second dark flash as a function of heart-rate.

**Figure S2.**
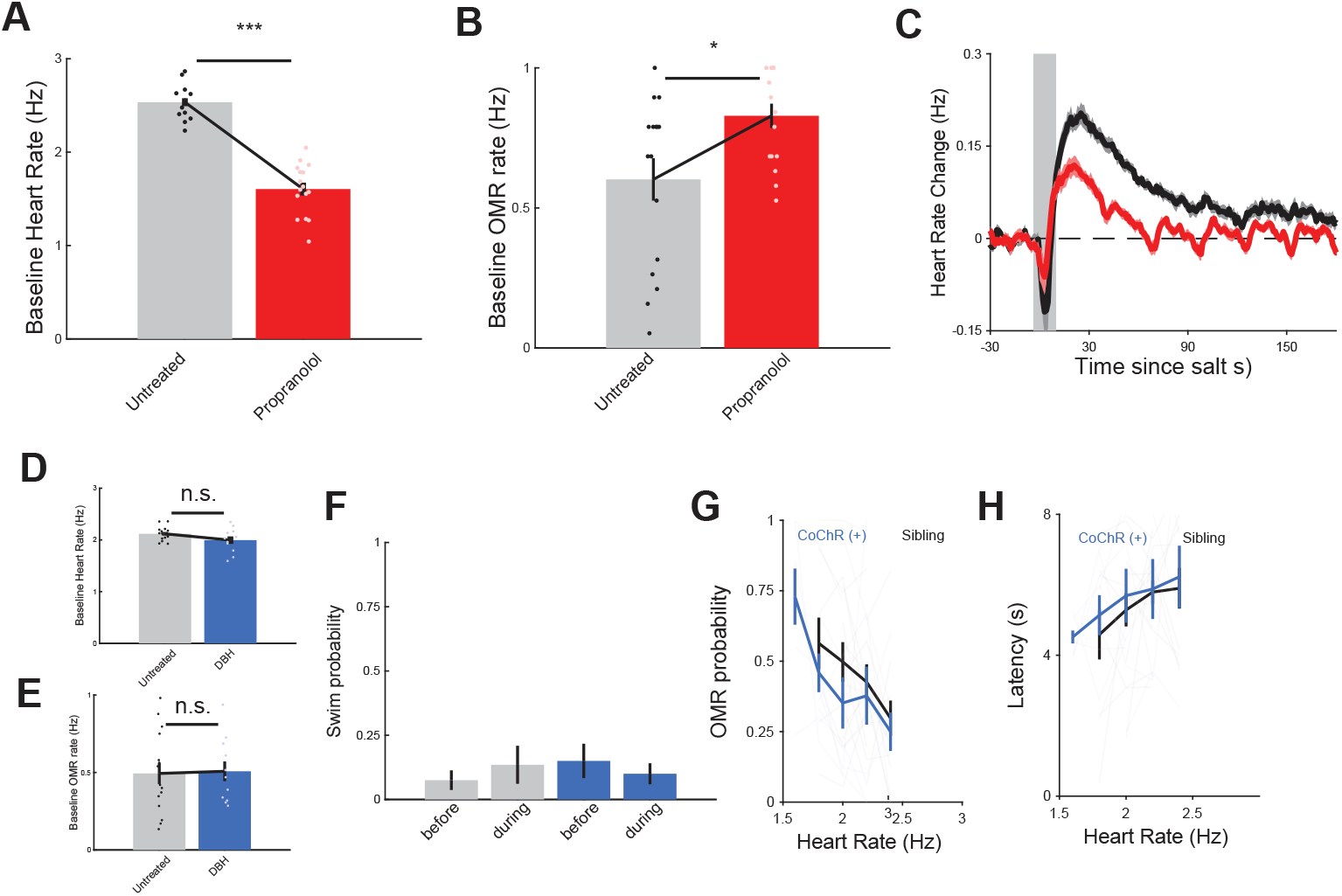
Supporting Material for Figure 2. (A) Baseline heart rate, from first 10 minutes of experiment, for larvae treated with propranolol (red) and sibling controls (gray). p = 1.54 *×* 10^−6^, wilcoxon rank sum. (B) Baseline optomotor response probability, from first 10 minutes of experiment, for larvae treated with propranolol (red) and sibling controls (gray). p = 0.0210, wilcoxon rank sum. (C) Stimulus-triggered heart rate for fish treated with propranolol (red) and sibling controls (black) following a salt pulse. (D) Baseline heart rate, from first 10 minutes of experiment, for *DβH:CoChR-eGFP* expressing larvae (blue) and sibling controls (gray). p = 0.1772, wilcoxon rank sum. (E) Baseline optomotor response rate, from first 10 minutes of experiment, for *DβH:CoChR-eGFP* expressing larvae (blue) and sibling controls (gray). p = 0.9173, wilcoxon rank sum. (F) Probability of a struggle during the 30 seconds before or 30 seconds during illumination with blue light for *DβH:CoChR-eGFP* expressing larvae (blue) and sibling controls (gray). (G) OMR probability as a function of heart rate for *DβH:CoChR-eGFP* expressing larvae (blue) and sibling controls (gray). (H) OMR response latency for *DβH:CoChR-eGFP* expressing larvae (blue) and sibling controls (gray).

**Figure S3.**
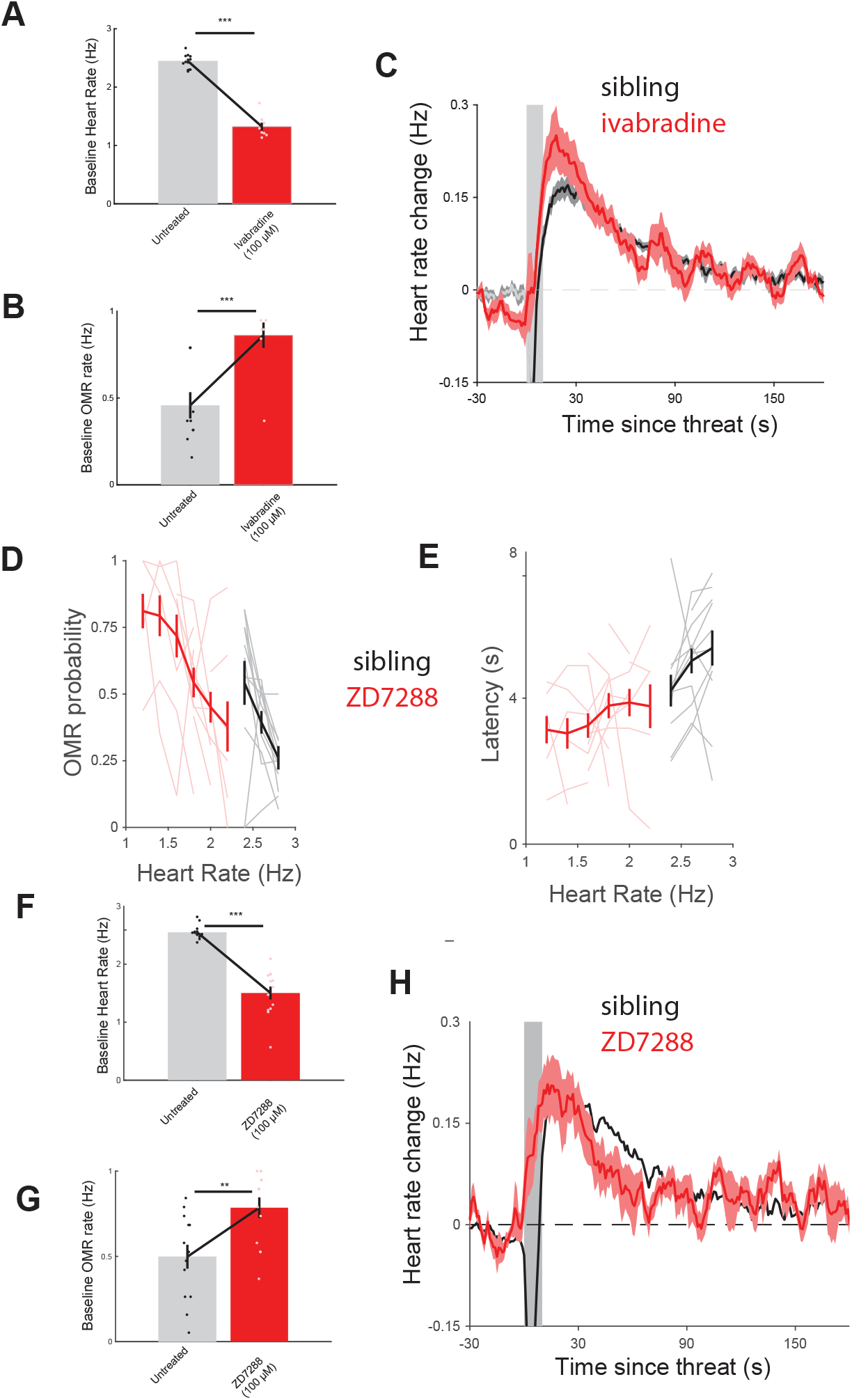
Supporting Material for Figure 3. (A) Baseline heart rate, from first 10 minutes of experiment, for larvae treated with ivabradine (red) and sibling controls (gray). p = 4.57*×* 10^−5^, wilcoxon rank sum. (B) Baseline optomotor response probability, from first 10 minutes of experiment, for larvae treated with ivabradine (red) and sibling controls (gray). p = 8.68 *×* 10^−4^, wilcoxon rank sum. (C) Stimulus-triggered heart rate for fish treated with ivabradine (red) and sibling controls (black) following a salt pulse. (D) OMR probability and (E) latency as a function of heart rate for fish treated with 100 *µ*M ZD7288 (red, n = 10) and sibling controls (gray, n =13) (F) Baseline heart rate, from first 10 minutes of experiment, for larvae treated with 100 *µ*M ZD7288 (red) and sibling controls (gray). p = 1.65*×* 10^−5^, wilcoxon rank sum. (G) Baseline optomotor response probability, from first 10 minutes of experiment, for larvae treated with 100 *µ*M ZD7288 (red) and sibling controls (gray). p = 0.0069, wilcoxon rank sum. (H) Stimulus triggered heart rate for fish treated with 100 *µ*M ZD7288 (red) and sibling controls (black) following a salt pulse.

**Figure S4.**
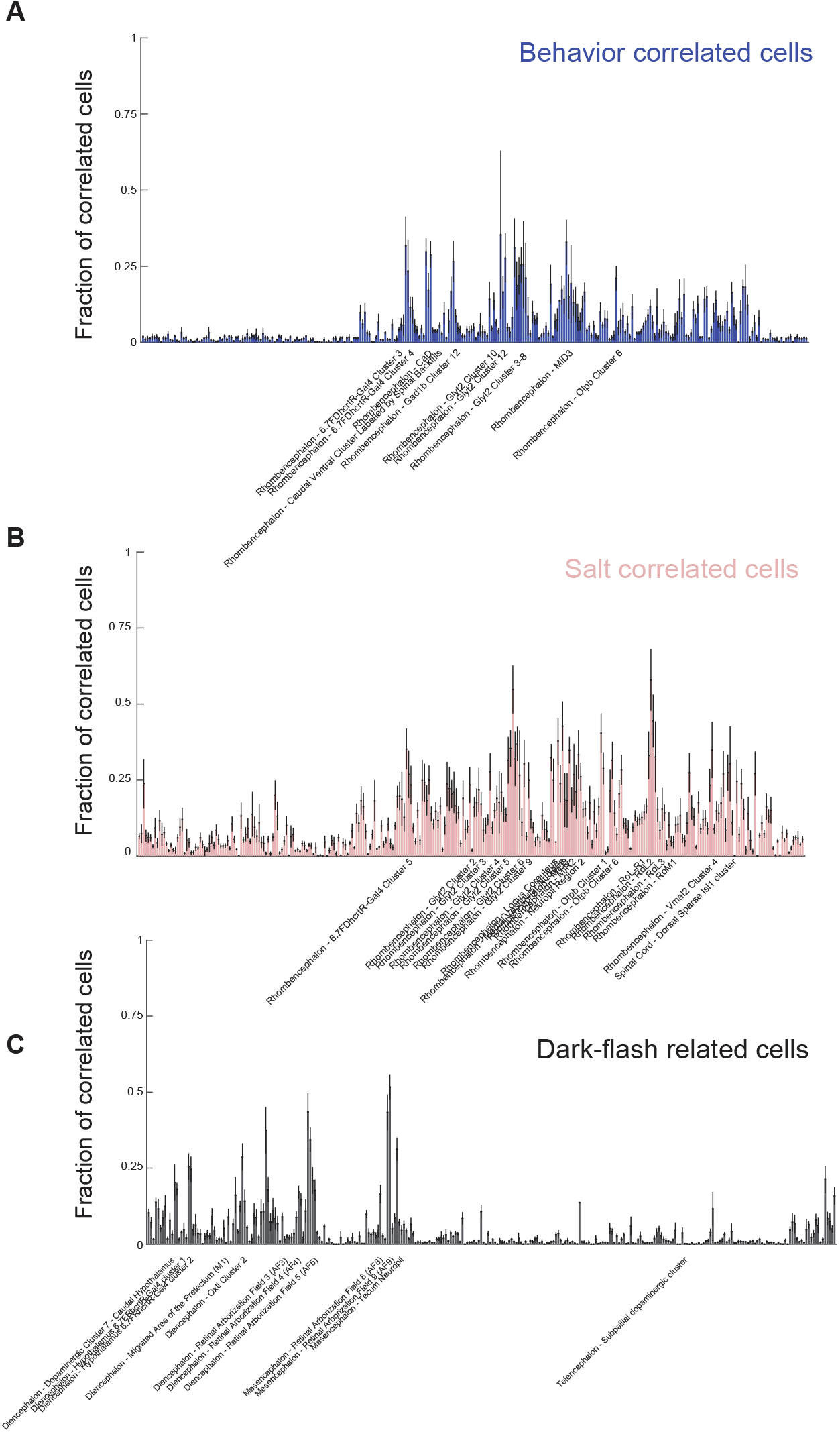
Supporting Material for Figure 5. (A) Fraction of cells in each Z-brain mask with strongest correlation to Behavior (masks with >20% are labeled). (B) Fraction of cells in each Z-brain mask with strongest correlation to salt pulse (masks with >30% are labeled). (C) Fraction of cells in each Z-brain mask with strongest correlation to salt pulse (masks with >20% are labeled).

## Notes

### Competing Interest Statement

The authors have declared no competing interest.

