## Supplemental Figures for "Cardiac interoception impacts behavior and brain-wide neuronal dynamics"

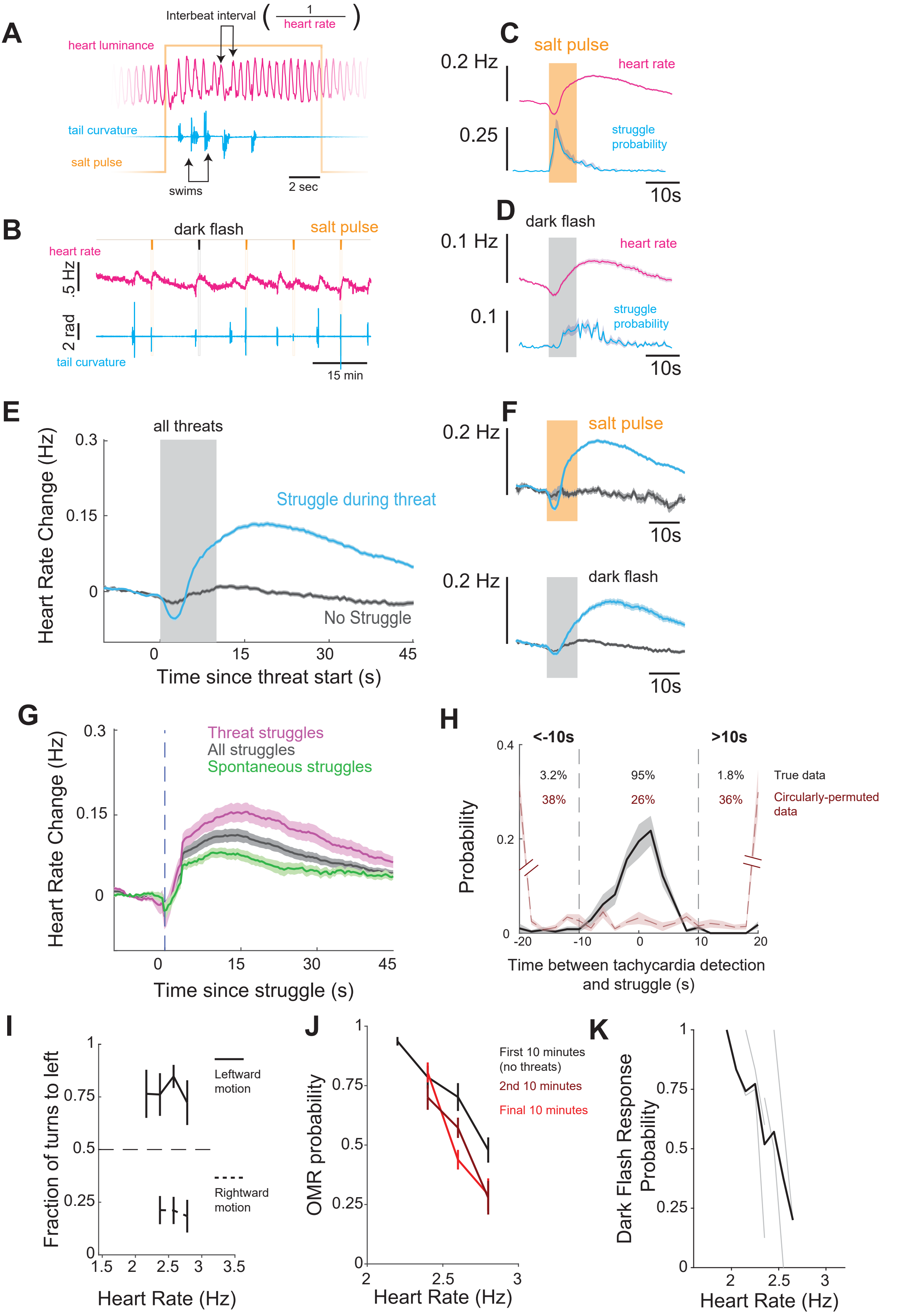

A
Interbeat interval
1
heart rate
heart luminance
tail curvature
salt pulse
swims
2 sec
C
salt pulse
0.2 Hz
heart rate
0.25
struggle probability
10s
B
dark flash
salt pulse
heart rate
.5 Hz
2 rad
tail curvature
15 min
D
dark flash
0.1 Hz
heart rate
0.1
struggle probability
10s
E
all threats
Heart Rate Change (Hz)
0.3
0.15
0
Struggle during threat
No Struggle
0
15
30
45
Time since threat start (s)
F
0.2 Hz
salt pulse
10s
0.2 Hz
dark flash
10s
G
Heart Rate Change (Hz)
0.3
0.15
0
Threat struggles
All struggles
Spontaneous struggles
0
15
30
45
Time since struggle (s)
H
<-10s
>10s
0.4
0.2
0
Probability
3.2%
95%
1.8%
True data
38%
26%
36%
Circularly-permuted data
-20
-10
0
10
20
Time between tachycardia detection and struggle (s)
I
Fraction of turns to left
1
0.75
0.5
0.25
0
Leftward motion
Rightward motion
1.5
2
2.5
3
3.5
Heart Rate (Hz)
J
OMR probability
1
0.75
0.5
0.25
0
First 10 minutes (no threats)
2nd 10 minutes
Final 10 minutes
2
2.5
3
Heart Rate (Hz)
K
Dark Flash Response Probability
1
0.75
0.5
0.25
0
2
2.5
3
Heart Rate (Hz)

**Figure S1. Supporting Material for Figure 1**
**(A)** Sample data from 12 seconds illustrating individual heart beats (pink) and tail events (blue) during a threat (salt, orange).
**(B)** Sample experiment showing different threats (darkflash, black, and salt pulse orange), heart rate (pink) and tail (blue) over 30 minutes.
**(C)** Stimulus triggered heart rate (pink) and tail response probability (blue) following a salt pulse.
**(D)** Stimulus-triggered heart rate and tail response probability (blue) following a dark flash.
**(E)** Stimulus triggered heart rate following threats (either dark flash or salt pulse) where the fish struggled during the threat (blue) and those where it did not (gray).
**(F)** Data in (E) split for salt pulses (top) and dark flashes (bottom).
**(G)** Struggle-triggered heart rate average for all struggles (gray), those that ocurred during threats (pink) and those that occurred spontaneously (green).
**(H)** Histogram of the onset of tachycardia events relative to the time of the nearest struggle. Events separated greater than or less than 20 or -20 seconds are accumulated in the largest and smallest bins.
**(I)** Probability of the first turn being toward the left as a function of heart rate (x-axis) and whether or not the motion stimulus is to the left (top, solid) or right (bottom, dashed)
**(J)** Probability of responding to OMR trial versus heart rate for data separated into 3 epochs - first 10 minutes (black), during which no salt-pulses are delivered, and the second (brown) and final (red) 10 minutes, during which salt pulses are delivered every 6 minutes.
**(K)** Probability of a response to a 10 second dark flash as a function of heart-rate.

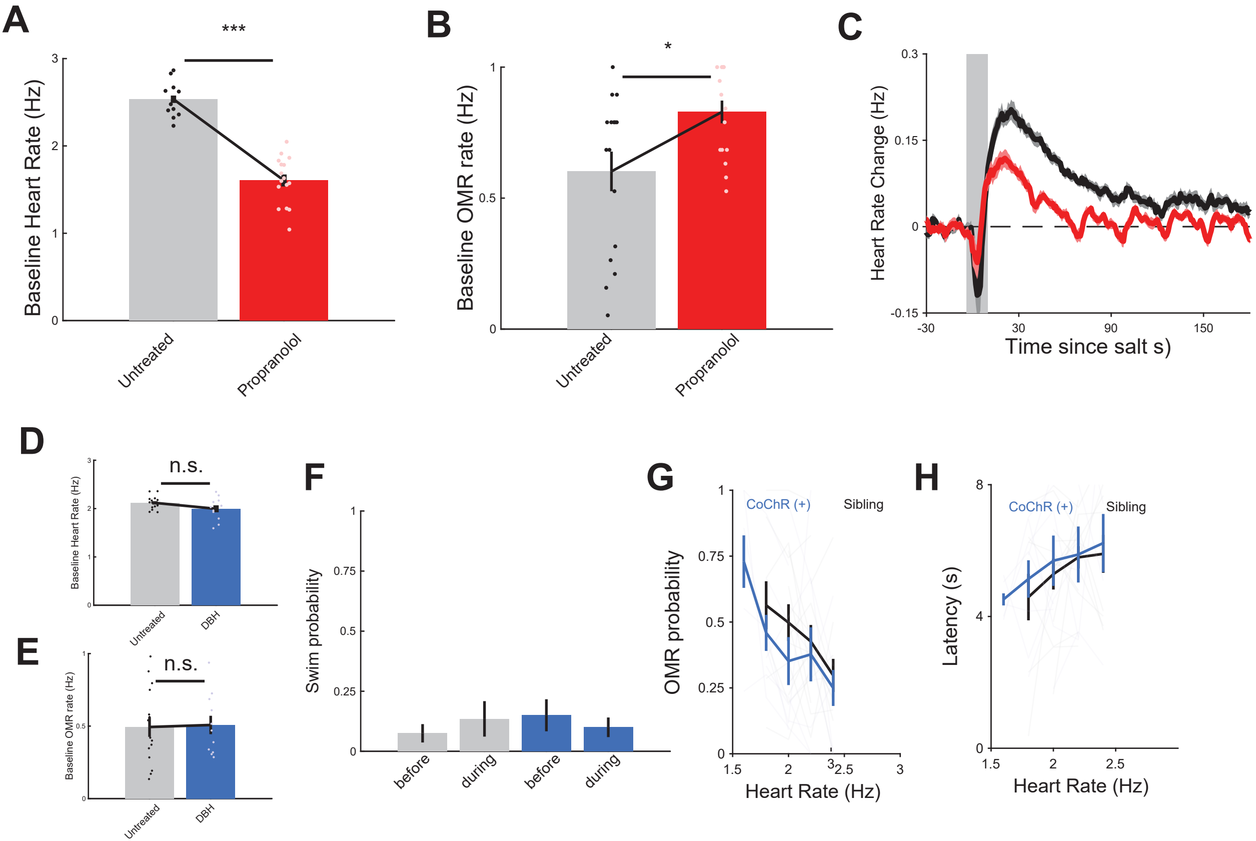

A
***
Baseline Heart Rate (Hz)
Untreated
Propranolol
B
*
Baseline OMR rate (Hz)
Untreated
Propranolol
C
Heart Rate Change (Hz)
Time since salt s)
D
n.s.
Baseline Heart Rate (Hz)
Untreated
DBH
E
n.s.
Baseline OMR rate (Hz)
Untreated
DBH
F
Swim probability
before
during
before
during
G
OMR probability
CoChR (+)
Sibling
Heart Rate (Hz)
H
Latency (s)
CoChR (+)
Sibling
Heart Rate (Hz)

**Figure S2. Supporting Material for Figure 2**

**(A)** Baseline heart rate, from first 10 minutes of experiment, for larvae treated with propranolol (red) and sibling controls (gray). p = 1.54×10–6, wilcoxon rank sum.

**(B)** Baseline optomotor response probability, from first 10 minutes of experiment, for larvae treated with propranolol (red) and sibling controls (gray). p = 0.0210, wilcoxon rank sum.

**(C)** Stimulus-triggered heart rate for fish treated with propranolol (red) and sibling controls (black) following a salt pulse.

**(D)** Baseline heart rate, from first 10 minutes of experiment, for DβH:CoChR-eGFP expressing larvae (blue) and sibling controls (gray). p = 0.1772, wilcoxon rank sum.

**(E)** Baseline optomotor response rate, from first 10 minutes of experiment, for DβH:CoChR-eGFP expressing larvae (blue) and sibling controls (gray). p = 0.9173, wilcoxon rank sum.

**(F)** Probability of a struggle during the 30 seconds before or 30 seconds during illumination with blue light for DβH:CoChR-eGFP expressing larvae (blue) and sibling controls (gray).

**(G)** OMR probability as a function of heart rate for DβH:CoChR-eGFP expressing larvae (blue) and sibling controls (gray).

**(H)** OMR response latency for DβH:CoChR-eGFP expressing larvae (blue) and sibling controls (gray).

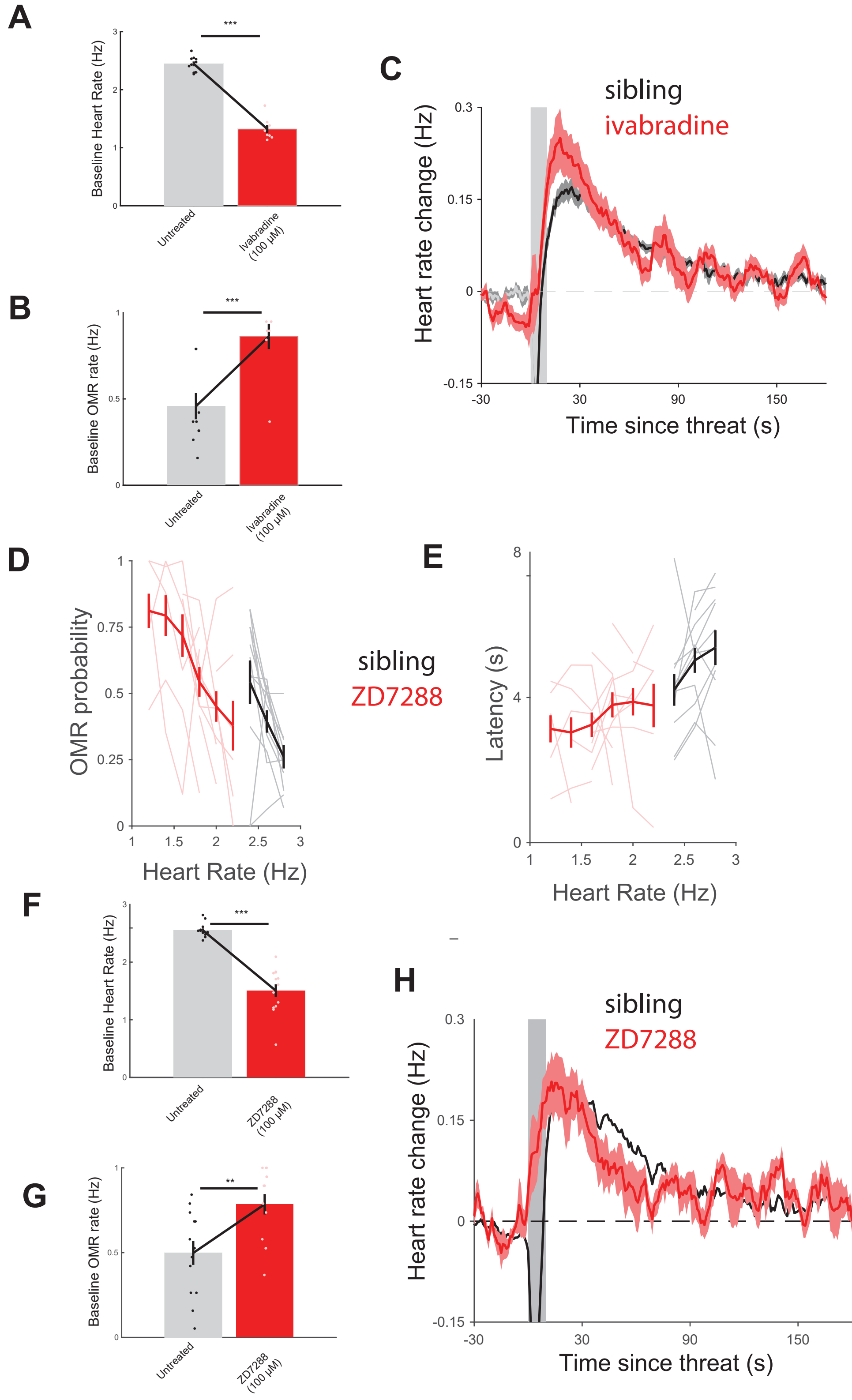
A
B
C
D
E
F
G
H
Baseline Heart Rate (Hz)
Untreated
Ivabradine (100 μM)
***
Baseline OMR rate (Hz)
sibling
ivabradine
Heart rate change (Hz)
Time since threat (s)
OMR probability
Heart Rate (Hz)
ZD7288
Latency (s)
ZD7288 (100 μM)
**

**Figure S3. Supporting Material for Figure 3**
**(A)** Baseline heart rate, from first 10 minutes of experiment, for larvae treated with ivabradine (red) and sibling controls (gray). p = 4.57 × 10–5, wilcoxon rank sum.
**(B)** Baseline optomotor response probability, from first 10 minutes of experiment, for larvae treated with ivabradine (red) and sibling controls (gray). p = 8.68 × 10–4, wilcoxon rank sum.
**(C)** Stimulus-triggered heart rate for fish treated with ivabradine (red) and sibling controls (black) following a salt pulse.
**(D)** OMR probability and **(E)** latency as a function of heart rate for fish treated with 100 μM ZD7288 (red, n = 10) and sibling controls (gray, n =13).
**(F)** Baseline heart rate, from first 10 minutes of experiment, for larvae treated with 100 μM ZD7288 (red) and sibling controls (gray). p = 1.65 × 10–5, wilcoxon rank sum.
**(G)** Baseline optomotor response probability, from first 10 minutes of experiment, for larvae treated with 100 μM ZD7288 (red) and sibling controls (gray). p = 0.0069, wilcoxon rank sum.
**(H)** Stimulus triggered heart rate for fish treated with 100 μM ZD7288 (red) and sibling controls (black) following a salt pulse.

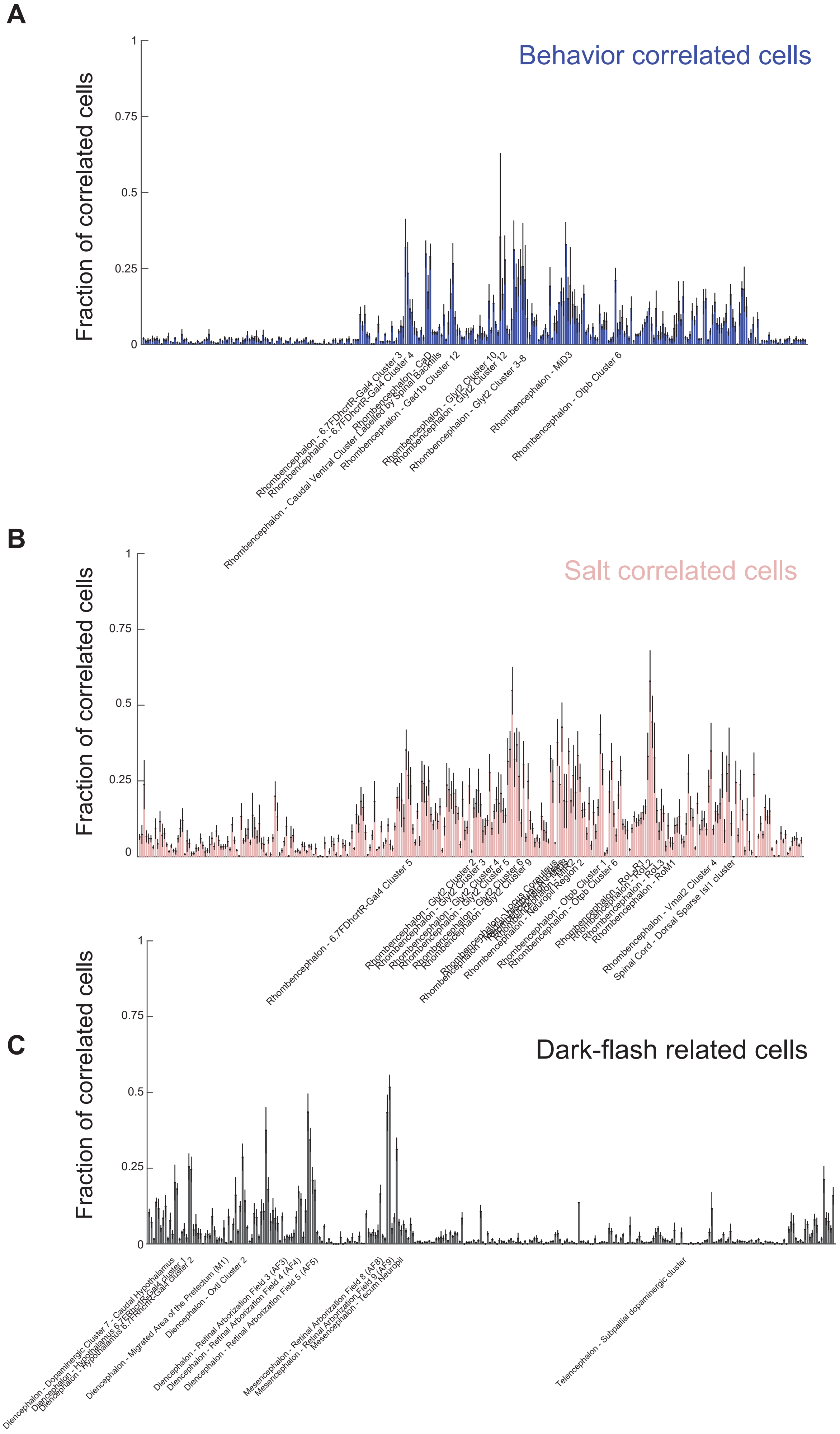
A
Behavior correlated cells
Fraction of correlated cells
1
0.75
0.5
0.25
0
Rhombencephalon - 6.7FDhcrtR-Gal4 Cluster 3
Rhombencephalon - 6.7FDhcrtR-Gal4 Cluster 4
Rhombencephalon - CaD
Rhombencephalon - Caudal Ventral Cluster Labelled by Spinal Backfills
Rhombencephalon - Gad1b Cluster 12
Rhombencephalon - Glyt2 Cluster 10
Rhombencephalon - Glyt2 Cluster 12
Rhombencephalon - Glyt2 Cluster 3-8
Rhombencephalon - MiD3
Rhombencephalon - Otpb Cluster 6
B
Salt correlated cells
Fraction of correlated cells
1
0.75
0.5
0.25
0
Rhombencephalon - 6.7FDhcrtR-Gal4 Cluster 5
Rhombencephalon - Glyt2 Cluster 2
Rhombencephalon - Glyt2 Cluster 3
Rhombencephalon - Glyt2 Cluster 4
Rhombencephalon - Glyt2 Cluster 5
Rhombencephalon - Glyt2 Cluster 6
Rhombencephalon - Glyt2 Cluster 9
Rhombencephalon - Locus Coreuleus
Rhombencephalon - Neuropil Region 2
Rhombencephalon - Otpb Cluster 1
Rhombencephalon - Otpb Cluster 6
Rhombencephalon - RoL-R1
Rhombencephalon - RoL2
Rhombencephalon - RoL3
Rhombencephalon - RoM1
Rhombencephalon - Vmat2 Cluster 4
Spinal Cord - Dorsal Sparse Isl1 cluster
C
Dark-flash related cells
Fraction of correlated cells
1
0.75
0.5
0.25
0
Diencephalon - Dopaminergic Cluster 7 - Caudal Hypothalamus
Diencephalon - Hypothalamus 6.7FDhcrtR-Gal4 cluster 1
Diencephalon - Hypothalamus 6.7FDhcrtR-Gal4 cluster 2
Diencephalon - Migrated Area of the Pretectum (M1)
Diencephalon - Oxtl Cluster 2
Diencephalon - Retinal Arborization Field 3 (AF3)
Diencephalon - Retinal Arborization Field 4 (AF4)
Diencephalon - Retinal Arborization Field 5 (AF5)
Mesencephalon - Retinal Arborization Field 8 (AF8)
Mesencephalon - Retinal Arborization Field 9 (AF9)
Mesencephalon - Tecum Neuropil
Telencephalon - Subpallial dopaminergic cluster

**Figure S4. Supporting Material for Figure 5**
**(A)** Fraction of cells in each Z-brain mask with strongest correlation to Behavior (masks with >20% are labeled).
**(B)** Fraction of cells in each Z-brain mask with strongest correlation to salt pulse (masks with >30% are labeled).
**(C)** Fraction of cells in each Z-brain mask with strongest correlation to salt pulse (masks with >20% are labeled).
